# Algorithmic biosynthesis of eukaryotic glycans

**DOI:** 10.1101/440792

**Authors:** Anjali Jaiman, Mukund Thattai

## Abstract

An algorithm converts inputs to corresponding unique outputs through a sequence of actions. Algorithms are used as metaphors for complex biological processes such as organismal development. Here we make this metaphor rigorous for glycan biosynthesis. Glycans are branched sugar oligomers that are attached to cell-surface proteins and convey cellular identity. Eukaryotic O-glycans are synthesized by collections of enzymes in Golgi compartments. A compartment can stochastically convert a single input oligomer to a heterogeneous set of possible output oligomers; yet a given type of protein is invariably associated with a narrow and reproducible glycan oligomer profile. Here we resolve this paradox by borrowing from the theory of algorithmic self-assembly. We rigorously enumerate the sources of glycan microheterogeneity: incomplete oligomers via early exit from the reaction compartment; tandem repeat oligomers via runaway reactions; and competing oligomer fates via divergent reactions. We demonstrate how to diagnose and eliminate each of these, thereby obtaining “algorithmic compartments” that convert inputs to corresponding unique outputs. Given an input and a target output we either prove that the output cannot be algorithmically synthesized from the input, or explicitly construct an ordered series of algorithmic compartments that achieves this synthesis. Our theoretical analysis allows us to infer the causes of non-algorithmic microheterogeneity and species-specific diversity in real glycan datasets.

---

**A**N algorithm converts distinct inputs (e.g. a list of *n* numbers) to corresponding unique outputs (e.g. the same numbers sorted in increasing order) through a sequence of deterministic or stochastic actions (e.g. swaps of numbers at two positions). Algorithms typically represent compressed descriptions of the output, encoding a recipe for it rather than a copy of it (e.g. a list is sorted by many successive swaps, not by looking up a table of pre-sorted lists) [1].

The process by which an egg is converted to an adult during organismal development is plausibly algorithmic [2]: distinct genomes produce distinct unique adults; and the genome encodes a recipe to make an adult, it is not a homunculus of the adult. In this spirit we define a biosynthetic system as implementing a non-trivial algorithm if: (a) it can accept many distinct inputs; (b) it converts each input to a corresponding unique output; and (c) this is achieved without requiring a template of the output. Enzymatic catalysis and ribosomal translation are trivial instances of algorithms: an enzyme that converts a specific substrate to a specific product has a single allowed input-output pair; ribosomes assemble input amino acids into distinct output proteins by essentially copying an mRNA template, not through a compressed recipe. Here we study what is arguably the simplest biomolecular system capable of implementing a non-trivial algorithm: the eukaryotic glycan biosynthetic apparatus.

Glycans are branched sugar oligomers covalently attached to proteins on the surfaces of all living cells [3]. Eukaryotic glycans are composed of a small set of monosaccharide building block types (monomers) and disaccharide bond types (linkages in the branched glycan oligomer) [4] (Fig. 1A), and are attached to specific amino acid sites on the substrate protein. A given site on a given type of protein is typically associated with multiple types of oligomers (a phenomenon known as microheterogeneity) that collectively comprise the protein’s glycan profile [3, Chapter 1]. There is wide glycan diversity across eukaryotes since the same monomers can be assembled in an astronomical array of oligomeric combinations [5], [6]. Nevertheless, the glycan profiles of individual proteins are typically narrow and reproducible [7]. These glycan profiles can vary between species [8]–[10]; between individuals [11], as with ABO blood groups [12]; and between cell types in an individual [13]. Species-specific glycan profiles facilitate self-nonself identification, while cell-specific glycan profiles have roles in development [14]–[16]. Narrow, reproducible glycan profiles are functionally relevant since glycan alterations are implicated in cancers and genetic diseases [17]–[21].

**Fig. 1.**
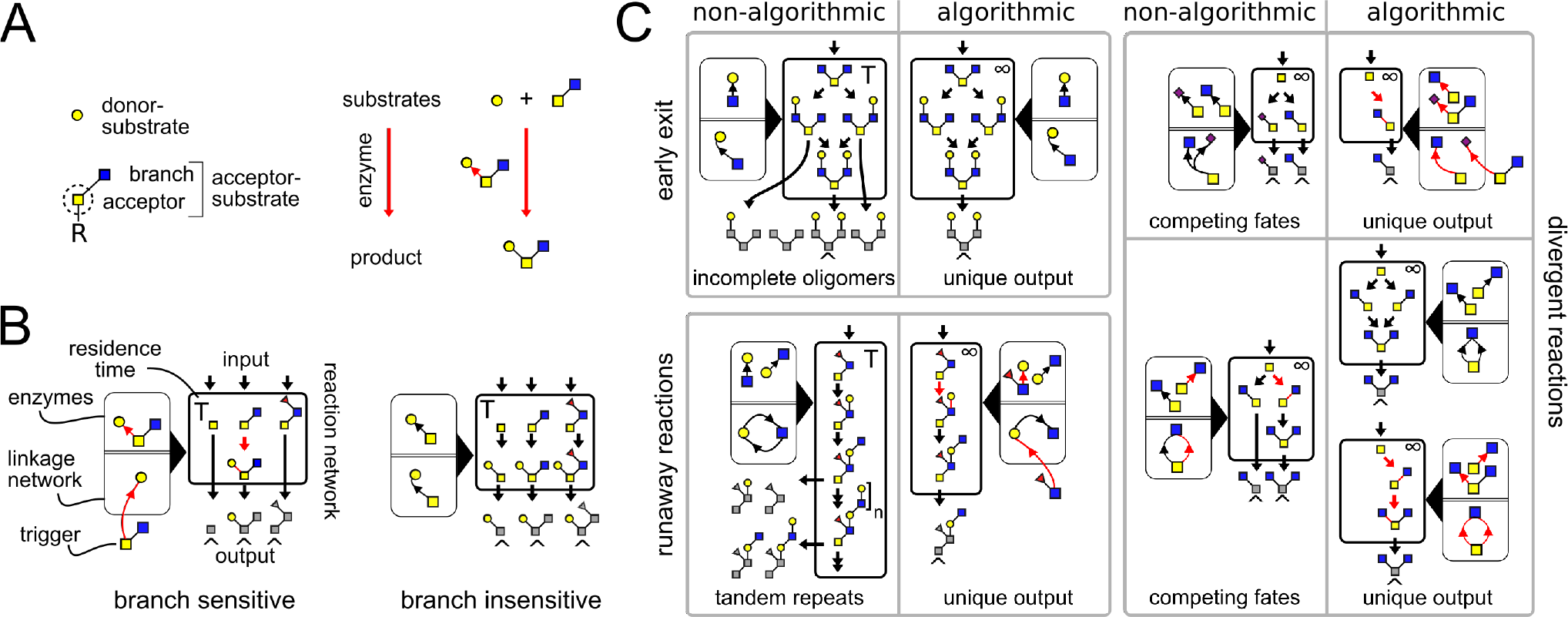
Glycan biosynthesis and microheterogeneity. *(A)* A GTase enzyme catalyzes a specific carbon-carbon linkage between a specific donor monomer type (the “donor-substrate”) and a specific acceptor monomer type with specific branches (the “acceptor-substrate”) on some arbitrary oligomer (“R”). We represent distinct monomer types in an oligomer by shapes/colors, and linkages between distinct monomer carbons by distinct bond angles [3]. We represent each GTase enzyme graphically, showing its acceptor-substrate and the specific monomer-addition reaction it catalyzes (arrow from acceptor to donor at distinct angles for distinct acceptor carbons). This notation makes it easy to identify acceptor-substrates in an input oligomer, and to predict the resulting output oligomer. *(B)* A compartment (bold box) contains GTase enzymes and converts input oligomers to output oligomers. In place of full input oligomers we often show only the acceptor-substrate of interest. Within the compartment we represent oligomer growth by a reaction network whose arrows show single-monomer-addition reactions. Oligomers exit the compartment as outputs after an average residence time T. We sometimes shade in gray the portions of the output oligomers provided as inputs. Terminal oligomers of a compartment (endpoints of the reaction network, labeled ^) cannot be further extended in that compartment. Next to the compartment (light box, black triangle) we show the enzymes it contains (above the double line) and their corresponding linkage network (below the double line). The linkage network (DEFINITIONS) represents all the orders in which monomer types can be linked, with arrows from acceptor monomer types to donor monomer types (at distinct angles for distinct acceptor carbons). Triggers (DEFINITIONS) are acceptor-substrates whose branches cannot be fully synthesized within the compartment; they must be provided as inputs to the compartment. Left: Branch-sensitive enzymes (red enzyme and reaction arrows) recognize as their acceptor-substrates only specific monomer types that have or lack specific branches. Right: Branch-insensitive enzymes (black enzyme and reaction arrows) recognize as their acceptor-substrates specific acceptor monomer types regardless of their branches. *(C)* We show three sources of glycan microheterogeneity (non-algorithmic, an input gives many possible outputs) and show how each of these may be eliminated (algorithmic, an input gives a unique output). “Early exit”: Oligomers could exit the compartment at a random intermediate stage of growth, before being converted to a terminal oligomer. This gives incomplete oligomers. Algorithmic solution: Ensure the compartment residence time is sufficiently long (labeled ∞) so all input oligomers are converted to terminal oligomers before exit. “Runaway reactions”: One or more enzymes could drive a runaway reaction (if and only if the compartment’s linkage network has a loop; Fig. 2A). This gives oligomers with an arbitrary number of tandem repeats. Algorithmic solution: Break the loop by ensuring some enzyme in the loop requires a trigger. “Divergent reactions”: Different enzymes could drive the same oligomer into divergent paths that never reconverge (only if the compartment has an acceptor conflict; Fig. 2B). This gives competing oligomer fates depending on the random order of enzyme action. Top-right: Two enzymes could act on the same acceptor-substrate in a mutually exclusive manner (e.g. at the same carbon). Algorithmic solution: Add branch sensitivity to ensure no two enzymes can act on the same acceptor-substrate. Bottom-right: The action of one enzyme could block the subsequent action of another. Algorithmic solution: either remove branch sensitivity so the enzymes can act in any order, or add further branch sensitivity so the enzymes must act in a fixed order.

Eukaryotic glycan oligomers are assembled by collections of glycosyltransferase (GTase) enzymes in the ER and Golgi apparatus, a process known as glycosylation. Here we focus on the class of eukaryotic O-glycans [3, Chapter 10], whose synthesis begins in the Golgi apparatus when a root monomer is attached to a serine or threonine on the substrate protein. The oligomer grows one monomer at a time as the protein moves through successive Golgi compartments, stochastically encountering various GTase enzymes within each compartment [22]. Different protein types encounter different subsets of GTase enzymes via sequence-specific interactions, and thus acquire distinct glycan profiles [3, Chapter 5,6]. How can this stochastic and heterogeneous biosynthetic process generate narrow and reproducible glycan profiles?

To resolve this paradox we borrow from the field of algorithmic self-assembly, which explores how small building blocks with stochastic local interactions assemble into various global configurations [23]–[25]. A central inverse problem in self-assembly is to design building blocks that assemble into a target shape. This process has been studied using the theoretical framework of Wang tiles: square building blocks with colored sides that stick to one another along sides with matching colors [26]. For efficient assembly of Wang tiles, their coloring must be cleverly chosen so the target shape is the unique terminal output of the stochastic assembly process [27]. That is, their assembly must be algorithmic. Glycans may be considered a natural realization of the Wang construct, with monomers acting like tiles whose stickiness is encoded by GTase enzymes. If glycan biosynthesis could be made algorithmic, cells could suppress unwanted byproducts and generate only desired glycan oligomers with high yield.

Here we determine the precise conditions under which glycan biosynthesis is algorithmic. We identify sources of microheterogeneity, and show how each of these may be eliminated. We then present a procedure to verify whether a desired target oligomer can be uniquely synthesized from a given input. This idealized algorithmic limit gives insight into the the biologically relevant case: glycan profiles having a few oligomers rather than a single unique oligomer. Our results complement systems-level analyses of glycosylation [7], [28]–[30], and provide a unified explanation for many enigmatic features of glycan profiles observed in nature. Moreover, the abstract mathematical structure at the heart of this remarkable biosynthetic system generalizes beyond specific biological details, and can therefore inform artificial algorithmic self-assembly efforts.

## RESULTS

### Glycan biosynthesis by GTase enzymes

A given GTase enzyme catalyzes a specific carbon-carbon linkage between a specific free donor monomer type and a specific acceptor monomer type on the growing oligomer (Fig. 1A; DEFINITIONS). Beyond this chemical specificity, GTase enzymes are often branch-sensitive (Fig. 1B, left): some act only if the acceptor monomer is empty, others only if the acceptor monomer is already linked to a specific branch [3, Chapter 6,10] [31], [32]. There is no strong evidence for more complex types of GTase enzyme specificity or proofreading in O-glycan synthesis. In particular, a GTase enzyme cannot distinguish between multiple instances of the same acceptor monomer (with the same branches) on an oligomer [22], [29]. We therefore define the acceptor-substrate of an idealized GTase enzyme to be a specific acceptor monomer type with its specific branches, rather than the entire oligomer (Fig. 1A). A real GTase enzyme might act on many different branched variants of an acceptor monomer (for example, it might only require that the first monomer of a branch be of a specific type); this is equivalent to a collection of idealized GTase enzymes, one for each possible acceptor-substrate. For the special case of a branch-insensitive GTase enzyme (Fig. 1B, right) we use the empty acceptor monomer as a shorthand for its many possible branched acceptor-substrates. We often refer to an acceptor-substrate itself as the input to a compartment, leaving implicit the larger oligomer of which it is a part.

### Sources of glycan microheterogeneity

Glycan microheterogeneity arises because oligomer growth is fundamentally stochastic. A compartment will contain a specific set of GTase enzymes responsible for growing an oligomer on a specific protein type. The order in which the growing oligomer encounters these enzymes is equivalent to a process of random sampling with replacement, with randomly distributed time intervals between successive encounters [33] (PROOFS: Remark 1). The reaction network (DEFINITIONS) of a compartment shows every possible oligomer growth order starting from a given input oligomer, as a result of all possible enzyme-catalyzed single-monomer-addition reactions in all possible permutations (Fig. 1B). Within a reaction network, incomplete oligomers are those that can potentially be further extended by some available GTase enzyme, and terminal oligomers are those that cannot be further extended. Two identical input oligomers might take different paths in the reaction network as they encounter GTase enzymes in different random permutations and at different random times, including the possibility of never encountering some enzyme or encountering the same enzyme repeatedly. Finally, each growing oligomer exits the compartment as an output after some average residence time *T*.

For compartments containing GTase enzymes with the properties discussed here, microheterogeneity can arise in precisely three ways (PROOFS: Remark 1). Two identical input oligomers might exit the compartment at different stages in the reaction network, giving a combination of both incomplete and terminal oligomers as outputs (“early exit”, Fig. 1C). The reaction network starting from some input oligomer might contain infinite runaway reactions, giving oligomers with arbitrary numbers of tandem repeats as outputs (“runaway reactions”, Fig. 1C,2A). Or the reaction network starting from some input oligomer might contain divergent reactions, giving competing oligomer fates as outputs (“divergent reactions”, Fig. 1C,2B). For a given compartment, it is possible that some input oligomers generate heterogeneous outputs, while others do not. Ideally we would like to assess the potential for microheterogeneity across all possible inputs. We now show how to diagnose each type of microheterogeneity from properties of the compartment as a whole, rather than having to track all reaction networks starting from all possible inputs. These results are summarized in Table 1.

**Fig. 2.**
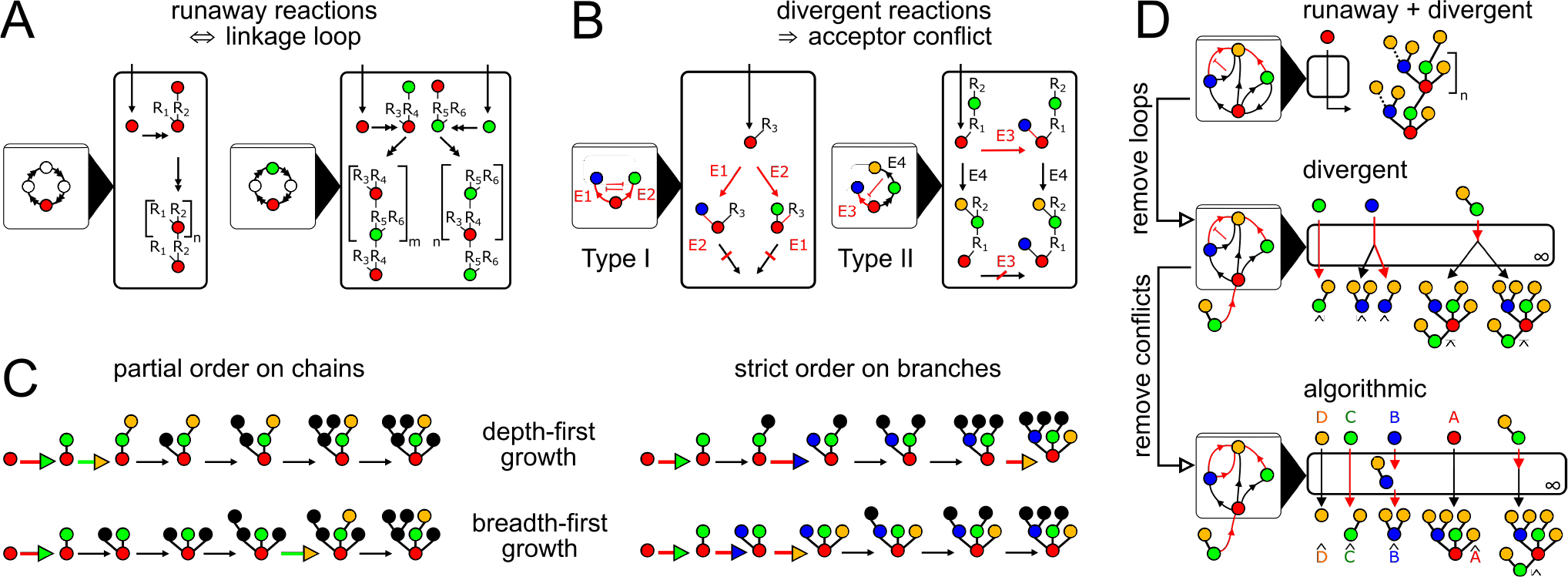
Algorithmic compartments and uniform depth-first growth. Colors represent monomer types (red, A; blue, B; green, C; orange, D); “R_i_” represents an arbitrary oligomer. Double arrows represent multiple reaction steps; blunt red arrows in the linkage network represent the inhibition of one enzyme by the action of another. ***(A)*** Runaway reactions occur whenever certain steps of oligomer growth can be iterated to produce tandem repeats. Loops in the linkage network are necessary and sufficient for runaway reactions (PROOFS: Lemma 1). For example: monomer A is added to a branch of monomer A, ad infinitum (left); monomer C is added to a branch of monomer A, and monomer A is added to a branch of monomer C, ad infinitum (right). ***(B)*** Divergent reactions occur whenever the reaction network has a fork that can never reconverge. This occurs when the action of one enzyme blocks the subsequent action of another, so the fate of the oligomer depends on the random order of enzyme action. Acceptor conflicts are necessary for divergent reactions (PROOFS: Lemma 2). Type I acceptor conflict: two enzymes compete for the same acceptor-substrate. This effect can be unidirectional or bidirectional. In this example E1 and E2 compete for the same acceptor-substrate; if E2 acts first then E1 is blocked from acting, and vice versa. Type II acceptor conflict: the acceptor-substrate of one enzyme is on some branch of the acceptor-substrate of another. This effect is always unidirectional. In this example E4 acts on a branch of the acceptor-substrate of E3; if E4 acts first then E3 is blocked from acting. ***(C)*** Chain order is the order in which different monomer types are added from root to tip along any chain. Branch order is the order in which different carbons of each acceptor-substrate are linked to donor monomers. Growth order is the order in which an oligomer grows, one monomer at a time. Two of the many possible growth orders are shown: a depth-first order (top row) and a breadth-first order (bottom row). A uniform depth-first growth order is defined as a depth-first growth order in which identical acceptor-substrates arising at any stage of growth are always linked to the same donor monomer type at the same carbon. This imposes a partial order on monomer types along chains, and a strict order of branch initiation on carbons of identical acceptor-substrate. ***(D)*** Suppose we are given a non-algorithmic compartment. We can eliminate early exit by ensuring an infinite residence time. This ensures only terminal oligomers are outputs. We can eliminate runaway reactions by removing or modifying at least one enzyme involved in each linkage loop. This imposes a partial order on chains, since every allowed chain corresponds to a directed walk along the acyclic linkage network. We can eliminate divergent reactions by removing all but one enzyme involved in each conflict. By eliminating Type I acceptor conflicts we ensure that no two enzymes have the same acceptor-substrate, imposing a strict order on branch additions for each acceptor-substrate. By eliminating Type II acceptor conflicts we ensure that all branches of all acceptor-substrates are terminal, imposing a depth-first growth order. The result is a conflict-free algorithmic compartment that converts each input oligomer to a corresponding unique output oligomer, via uniform depth-first growth. Any input-to-output map that can be achieved in any algorithmic compartment can also be achieved in such a conflict-free algorithmic compartment (PROOFS: Lemma 3).

**TABLE 1.** Sources of Glycan Microheterogeneity.

| Type of microheterogeneity | Reaction network property | Compartment property |
| --- | --- | --- |
| incomplete oligomers | ⇔ early exit | ⇔ finite residence |
| tandem repeats | ⇔ runaway reaction | ⇔ linkage loop |
| competing fates | ⇔ divergent reaction | ⇒ acceptor conflict |

*Early exit* will occur if and only if the compartment has a finite residence time, giving incomplete oligomers as outputs (PROOFS: Remark 1). Since there is no proofreading mechanism regulating oligomer exit, only at sufficiently long residence times (labeled *T* → ∞) will all input oligomers reach a terminal state before exit.

A *runaway reaction* is an infinite path in the reaction network, giving oligomers with arbitrary numbers of tandem repeats as outputs. To diagnose runaway reactions we must examine the compartment’s linkage network (Fig. 1B; DEFINITIONS), which shows all the orders in which monomer types may be linked to one another through the action of the available GTase enzymes. Certain GTase enzymes require triggers (branched acceptor-substrates that cannot be synthesized within the compartment itself; Fig. 1B; DEFINITIONS); these effectively act as novel monomer types in the linkage network. A compartment contains a runaway reaction if and only if its linkage network contains one or more loops (Fig. 2A; PROOFS: Lemma 1). Each such loop shows an order of monomer linkages that can be iterated to produce tandem repeats.

A *divergent reaction* is a fork in the reaction network that never reconverges, with distinct paths leading to competing oligomer fates as outputs. To diagnose divergent reactions we must examine the acceptor-substrates of every GTase enzyme in the compartment. A fork occurs whenever distinct enzymes can act on the same oligomer to yield distinct products. If these enzymes could act in any order (for example, if they act on distinct empty acceptor monomers on the oligomer) then the reaction paths could reconverge after the fork. A divergent reaction implies that the action of one enzyme on an oligomer blocks the subsequent action of another, so the reaction paths will never reconverge (Fig. 2B; PROOFS: Lemma 2): either one enzyme acts directly on the acceptor-substrate of another (Type I acceptor conflict; Fig. 2B, left) or one enzyme acts on a branch of the acceptor-substrate of another (Type II acceptor conflict; Fig. 2B, right) (DEFINITIONS). Note that the converse is not true: not all acceptor conflicts lead to divergent reactions, since the compartment might contain some other enzyme that can carry out the blocked reaction on the modified substrate.

### Algorithmic compartments

An algorithmic compartment is defined as one that converts each possible input oligomer to a corresponding unique output oligomer. We say the compartment achieves a well-defined input-to-output map. This means that every reaction network starting from every possible input converges to a corresponding unique terminal oligomer, and only these terminal oligomers exit as outputs. For a compartment to be algorithmic it is therefore necessary and sufficient to eliminate all three sources of microheterogeneity: early exit, runaway reactions, and divergent reactions (Fig. 2D; PROOFS: Remark 1). An algorithmic compartment must have infinite residence time (to eliminate early exit) and no linkage loops (to eliminate runaway reactions). Algorithmic compartments could have acceptor conflicts, since these do not always cause divergent reactions. However it is always possible to eliminate every acceptor conflict from an algorithmic compartment (by removing conflicted enzymes and thus eliminating certain growth orders; see PROOFS: Lemma 3) without changing the terminal oligomer of any reaction network. Conflicted enzymes are thus redundant: any input-to-output map that can be achieved in any algorithmic compartment can be achieved without acceptor conflicts.

### Uniform depth-first growth

We next define the key concept of uniform depth-first growth. There are many permutations in which a series of monomers may be added to a given initial oligomer to yield a given final oligomer. Among these possible growth orders, uniform depth-first growth orders (DEFINITIONS) are particularly relevant for analyzing glycan biosynthesis. “Depth-first” means no new branch is initiated on an acceptor monomer until all its existing branches are extended to their terminal fates (as set by the final oligomer). “Uniform” means, for identical acceptor monomers with identical terminally-extended branches, the new branch (if any) is always initiated at the same empty carbon by linking the same donor monomer type.

Under uniform depth-first growth, identical acceptor-substrates arising at any stage of growth will necessarily have the same fate in the final oligomer, and the same order in which their empty carbons are linked to new branches. The task of specifying a uniform depth-first growth order from a given initial oligomer to a given final oligomer is essentially reduced to specifying a consistent branch order (Fig. 2C, right). Note this does not completely fix the time order of growth. This is because an oligomer at an intermediate stage of growth could have many acceptor monomers with terminally-extended branches, each ready for a new branch to be initiated. These new branches could be added simultaneously, or in any permutation in time, without violating the uniform depth-first property.

Some initial/final oligomer pairs admit no uniform depth-first growth orders. This occurs, for example, if two identical acceptor-substrates in the initial oligomer have distinct fates in the final oligomer; or, if an acceptor-substrate that is extended at an intermediate stage of growth is present but unextended in the final oligomer. In particular, the final oligomer of a uniform depth-first growth order cannot include a tandem repeat cut off at an arbitrary length. This imposes a partial order on the addition of monomer types (*A* can’t be added twice along any chain; and if *B* follows *A* on some chain, then *A* can’t follow *B* on any chain; Fig. 2C, left).

Uniform depth-first growth is relevant because it fully captures what happens in an algorithmic compartment with infinite residence time, no linkage loops, and no acceptor conflicts (PROOFS: Lemma 3). Since the compartment is algorithmic, the reaction network starting from a given input oligomer has a single terminal oligomer. Since the residence time is infinite, every growth order starting from the input oligomer reaches this terminal oligomer as its final state. Since there are no Type II acceptor conflicts, growth is depth-first. And since there are no Type I acceptor conflicts, growth is uniform. Therefore every growth order starting from any given input oligomer in such a conflict-free algorithmic compartment is uniform depth-first (Fig. 2D).

### Algorithmic biosynthesis in a single compartment

We are now in a position to address the central inverse problem of algorithmic biosynthesis: how to convert a given input oligomer to a given target oligomer. As a first pass we could pick an arbitrary growth order and load a compartment with GTase enzymes corresponding to each successive single-monomer-addition reaction. This guarantees that the target oligomer will be synthesized from the given input. The trouble is, various other oligomers might also be synthesized due to early exit, runaway reactions and divergent reactions; and the target oligomer might itself be further extended at long residence times. For high yield without fine-tuning or proofreading, the input-to-target reaction network must have the target as its unique terminal oligomer. If this can be done, we say the input/target pair is algorithmically achievable. We show (Fig 3; PROOFS):

#### Theorem 1.

An input/target oligomer pair is algorithmically achievable in a single compartment if and only if there is a uniform depth-first growth order from the input to the target.

Intuitively, if a uniform depth-first growth order can be found, then a compartment loaded with the corresponding GTase enzymes will be algorithmic and we need not worry about runaway or divergent reactions. For the converse, recall that any input-to-output map that can be achieved in an algorithmic compartment can also be achieved in a conflict-free algorithmic compartment. We know that oligomer growth in a conflict-free algorithmic compartment must be uniform depth-first. Therefore if such a growth order cannot be found, then the target cannot be uniquely synthesized in any algorithmic compartment. Note that in an algorithmic compartment with acceptor conflicts, every reaction network will contain at least one growth order that is uniform depth-first (PROOFS: Lemma 3), but may also contain other growth orders that are not. For this reason, the uniform depth-first test only establishes that an input/target pair is algorithmically achievable; it does not follow that the actual oligomer growth order is uniform depth-first.

Our intuitive argument assumes the solution to the unique biosynthesis problem requires algorithmic compartments. What if there is some non-algorithmic compartment that uniquely converts the input to the target? The proof of Theorem 1 shows we need not consider this possibility: if there is any compartment that converts the input uniquely to the target, then there is guaranteed to be an algorithmic compartment that does the same thing. Algorithmic compartments represent a small subset of all possible compartments, but they are completely sufficient for unique biosynthesis.

Operationally, Theorem 1 converts the problem of determining whether an input/target pair is algorithmically achievable to the task of searching over possible growth orders. This is not a brute-force search, since we are interested only in the uniform depth-first property rather than the specific time order of growth. Therefore the problem is reduced to an efficient search over consistent branch orders. If no uniform depth-first growth order can be found, the input/target oligomer pair is not algorithmically achievable in a single compartment; such a case is examined in Fig. 3. We encourage the reader to work through this instructive example in detail, as it involves many of the core concepts of algorithmic biosynthesis.

**Fig. 3.**
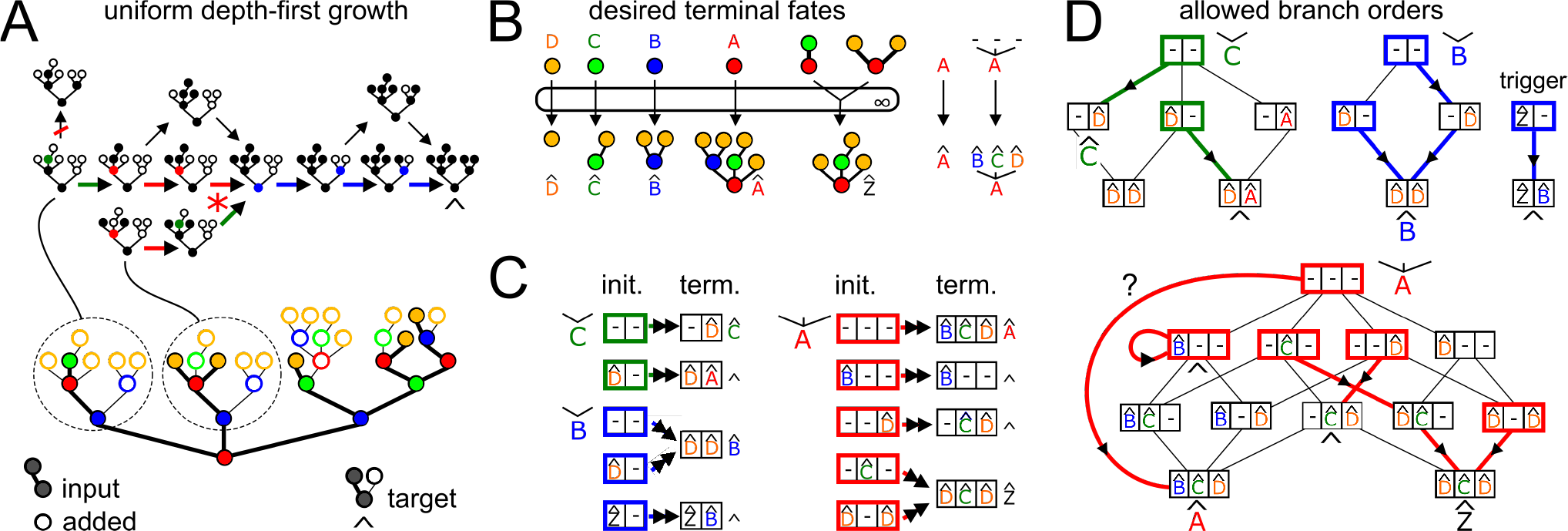
Algorithmic biosynthesis in a single compartment. ***(A)*** We ask whether a given input oligomer (filled circles, thick lines) can be uniquely converted to a given target oligomer (full structure) by the enzyme-catalyzed addition of new monomers (empty circles). By Theorem 1 this is possible if and only if we can find a uniform depth-first growth order from the input to the target. Top: Examples of possible growth orders for two sub-oligomers. At each step of growth an empty circle becomes filled as a new monomer is added. Depth-first growth means no new branch can be initiated on a monomer with incomplete existing branches; all reaction arrows shown here are consistent with depth-first growth, except for the leftmost vertical arrow. Uniform growth means identical acceptor-substrates are always linked to the same donor monomer type at the same empty carbon; thus if two growth paths converge to the same acceptor-substrate (red star) they must have the same subsequent growth step and the same terminal fate in the target oligomer (^). Alternative choices of branch order for a given acceptor-substrate correspond to alternative uniform depth-first growth orders (bifurcating arrows). The thick arrows show a depth-first growth order consistent with a particular choice of branch order; arrow colors show the monomer type of the growing acceptor-substrate. Note that the same branch order might be consistent with multiple growth orders (not shown here) since acceptor-substrates without ancestor-descendant relationships can grow independently. ***(B)*** Under uniform depth-first growth identical acceptor-substrates must have identical terminal fates in the target oligomer. The input oligomer contains many distinct acceptor-substrates, each potentially in multiple copies, and each with a desired terminal fate (labeled with a ^). A few examples of these are shown here. The terminal fates of empty monomers (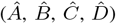) are of particular interest, since these comprise the allowed fates of any newly-initiated branches. The oligomer 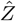 is a trigger branch: it cannot be fully synthesized starting from empty monomer *A.* Instead of the graphical representation (left) we use a recursive representation (right) showing the branches linked to each carbon of each acceptor-monomer type. Thus oligomer 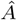 is monomer *A* linked to branches 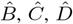 on its three carbons; oligomer 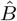 is monomer *B* linked to branch 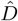 on both its carbons; oligomer 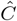 is monomer *C* linked to 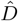 on its right carbon; and oligomer 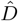 is simply monomer D. ***(C)*** The set of distinct initial acceptor-substrates (bold boxes colored by acceptor monomer type) and their desired terminal fates in the target oligomer (^). Under depth-first growth it is sufficient to consider how new branches are initiated on acceptor-substrates whose existing branches are already terminally extended. Boxes are colored according to acceptor monomer type; each slot shows any existing terminally-extended branches; empty carbons are labeled ‘–’. We have not shown the trivial case of monomer type D. ***(D)*** A uniform depth-first growth order is essentially determined by the choice of branch order. Depth-first growth is automatically enforced since we consider only acceptor-substrates with terminally-extended existing branches. Uniform growth requires that each distinct acceptor-substrate has a strict order on branch additions. We must find a single consistent branch order for each monomer type, such that each distinct acceptor-substrate achieves its desired terminal fate. The initiation of all possible branches in all possible orders is represented as a transition graph; we need consider only branches that are actually observed in the target oligomer. Each node of the graph represents distinct acceptor-substrates, using the box notation from Fig. 3C. Each directed edge represents the initiation and terminal extension of a branch on an empty carbon. Bold colored arrows show a possible choice of successive branch additions, from initial acceptor-substrates (bold boxes) to desired terminal fates (^). There can be no bold outward arrows from terminal fates; the bold self-loop represents an acceptor-substrate that is already at its desired terminal fate. Under uniform growth there can be only one bold outward arrow from each intermediate state. The branch order search might have a single unique solution, as with monomer type *C*; or multiple solutions, as with monomer type *B* (only one solution is shown here). There may be no solutions, since each choice of branch initiation cuts off other paths. In this example there is no set of bold arrows that simultaneously achieves all desired terminal fates for monomer type A. For the choice of arrows shown here, all desired terminal fates are achieved except for that of the empty monomer *A*. Therefore the given input/target pair is not algorithmically achievable in a single compartment.

### Algorithmic biosynthesis in multiple compartments

The Golgi apparatus consists of an ordered series of compartments containing distinct sets of GTase enzymes. Growing oligomers spend some average residence time in each successive enzymatic environment, reminiscent of a factory production line [22]. This is either because oligomers move through successive compartment types, or because the state of the compartment itself matures through successive enzymatic types [22], [34]. In this way, every output oligomer of each compartment type becomes an input for the next compartment type. How does the biosynthetic capacity of a multi-compartment series compare with that of a single compartment? We show (Fig. 4A; PROOFS):

#### Theorem 2.

An input/target oligomer pair is algorithmically achievable in a series of N compartments if and only if there is a growth order from the input to the target that can be fully decomposed into N uniform depth-first stretches.

The set of algorithmically achievable input/target pairs is evidently larger for a multi-compartment series than for a single compartment. We leave it as an exercise for the reader to show that the input/target pair from Fig. 3 is algorithmically achievable in two compartments via the use of triggers.

This result again highlights the power of algorithmic compartments. We might imagine that allowing a series to contain intermediate non-algorithmic compartments with multiple out-put oligomers might increase the number of input/target pairs that are algorithmically achievable. The proof of Theorem 2 shows this to be false: any unique synthesis that can be achieved with non-algorithmic compartments in series can always be achieved with the same number of algorithmic compartments in series.

Even allowing for multiple compartments, not all input/target pairs are algorithmically achievable. We show (Fig. 4B,C; PROOFS):

#### Corollary 2.1.

An input/target oligomer pair is algorithmically achievable if and only if there is a series of single-enzyme infinite-residence-time compartments that converts the input to the target as the unique final output.

This final result provides an efficient protocol to test whether an input oligomer can be uniquely converted to a target oligomer in any number of compartments. Moreover, it allows the construction of an explicit set of viable growth orders via enzyme-catalyzed single-monomer-addition reactions. Among these, the growth order with the fewest number of uniform depth-first stretches reveals the smallest number of compartments in which the unique synthesis of the target from the input can be achieved.

**Fig. 4.**
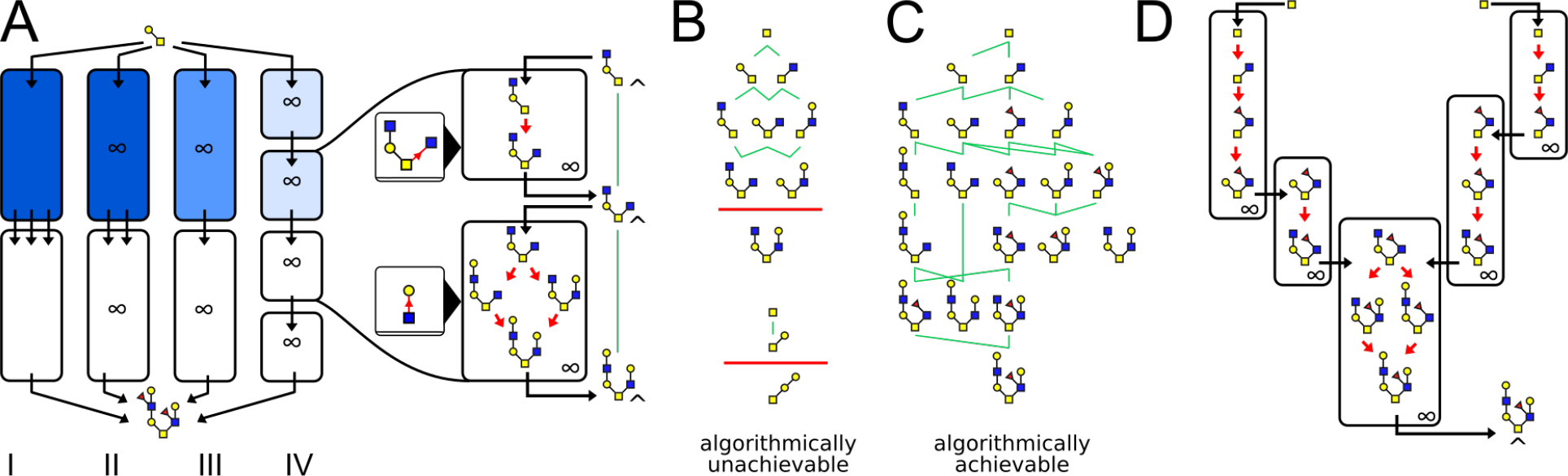
Algorithmic biosynthesis in multiple compartments. ***(A)*** Suppose we are given a series of compartments, not necessarily algorithmic, that convert a given input oligomer uniquely to a given target oligomer. We now proceed to modify the original compartments through successive steps (PROOFS: Theorem 2; the fading shade of blue represents the removal of enzymes in successive steps). We know none of the compartments contains a runaway reaction, since these would produce oligomers with arbitrary numbers of tandem repeats. By assumption every possible growth order through the reaction networks of every compartment in the series leads uniquely to the target oligomer. Step *I → II*: Assign every compartment an infinite residence time to eliminate incomplete oligomers. The resulting growth orders go via the terminal oligomers of each compartment and still lead to the target. Step *II → III*: Remove all enzymes with acceptor conflicts. This ensures each compartment is algorithmic while retaining at least one of the output oligomers of the corresponding original compartment, so all growth orders still lead uniquely to the target. Step *III → IV*: There is some growth order in which any given GTase enzyme acts successively on all its acceptor-substrates before another enzyme acts. This growth order persists when the original compartment is replaced with a series of infinite-residence-time compartments each containing a single GTase enzyme (square compartments). We represent the input-to-output map of each such compartment by a green edge. This construction shows that if an input/target pair is algorithmically achievable in a series of compartments, then it is achievable in a series of single-enzyme compartments. ***(B,C)*** We can check if an input-target pair is algorithmically achievable by searching for a series of infinite-residence-time single-enzyme compartments that convert the input (top) uniquely to the target (bottom). For each input/target pair we show all possible intermediate oligomers (sub-trees of the target and super-trees of the input). We connect two oligomers by a green edge if some infinite-residence-time single-enzyme compartment can uniquely convert the top oligomer to the bottom oligomer. An input/target oligomer pair is algorithmically achievable if and only if there is a continuous path via green edges from the input to the target. ***(B)*** Two target oligomers that cannot be uniquely synthesized from the input monomer. Top: Each GTase enzyme that could convert one of the penultimate oligomers to the target has multiple acceptor-substrates on the penultimate oligomer; the target is therefore an incomplete oligomer that will be further extended at infinite residence time. Bottom: The single-enzyme compartment required to convert the penultimate oligomer to the target contains a linkage loop; the target will therefore be further extended to form a tandem repeat. ***(C)*** This input-target pair is algorithmically achievable. There are seven distinct paths leading from the input to the target via single-enzyme compartments. ***(D)*** We can combine successive single-enzyme compartments as long as the corresponding growth order does not violate the uniform depth-first growth property. By carrying out this procedure over all possible paths from Fig. 4C we can find the minimum number of algorithmic compartments required to convert the input uniquely to the target (PROOFS: Theorem 2). Here we show two possible compartmental organizations both corresponding to the direct vertical path from the input to the target in Fig. 4C. Three compartments is the minimum number needed to uniquely synthesize the target oligomer.

## DISCUSSION

Algorithmic self-assembly has two basic conditions [27]: first, ensure that all growth paths from the input state lead to the target state and no further; second, wait long enough so the growing structure actually reaches the target state. All the complexity lies in the details of how to achieve the first condition; the second requires only patience. Our analysis begins by rigorously enumerating the sources of glycan microheterogeneity. We show that runaway and divergent reactions pull growth paths away from the desired target; by eliminating them we achieve the first condition. Early exit causes the production of incomplete structures; at sufficiently long residence times we achieve the second condition.

The actual algorithm that controls the fate of each input oligomer is encoded by the localization of GTase enzymes within the compartments of the Golgi apparatus. It has been argued that such enzyme compartmentalization allows for more controlled biosynthesis [22]. Our analysis reveals that the benefits of compartmentalization arise through multiple distinct mechanisms. First, by distributing enzymes across many compartments two sources of microheterogeneity – linkage loops and acceptor conflicts – are mitigated. Second, enzymes in earlier compartments can be used to construct triggers, exploited by enzymes in later compartments to achieve controlled oligomer growth.

To what extent is our algorithmic biosynthesis framework predictive? Given an oligomer we can infer the possible growth orders by which it could have been uniquely synthesized (Fig 3,4). We can also place a lower bound on the number of Golgi compartments required to achieve this synthesis (Fig. 4). In practice these results depend on having GTase enzymes with the required chemical specificity and branch sensitivity. Also, cells typically do not operate at the algorithmic limit: there is rarely just one unique oligomer associated with a given glycosylated protein. However, it is clear that real glycan profiles are astonishingly narrow compared to the potential biochemical diversity of sugar oligomers. By understanding how to synthesize a unique oligomer we can learn a great deal about how to synthesize a few related oligomers. In particular, we can understand which of the three distinct sources of microheterogeneity explain the observed glycan profile of real proteins (Fig. 5).

**Fig. 5.**
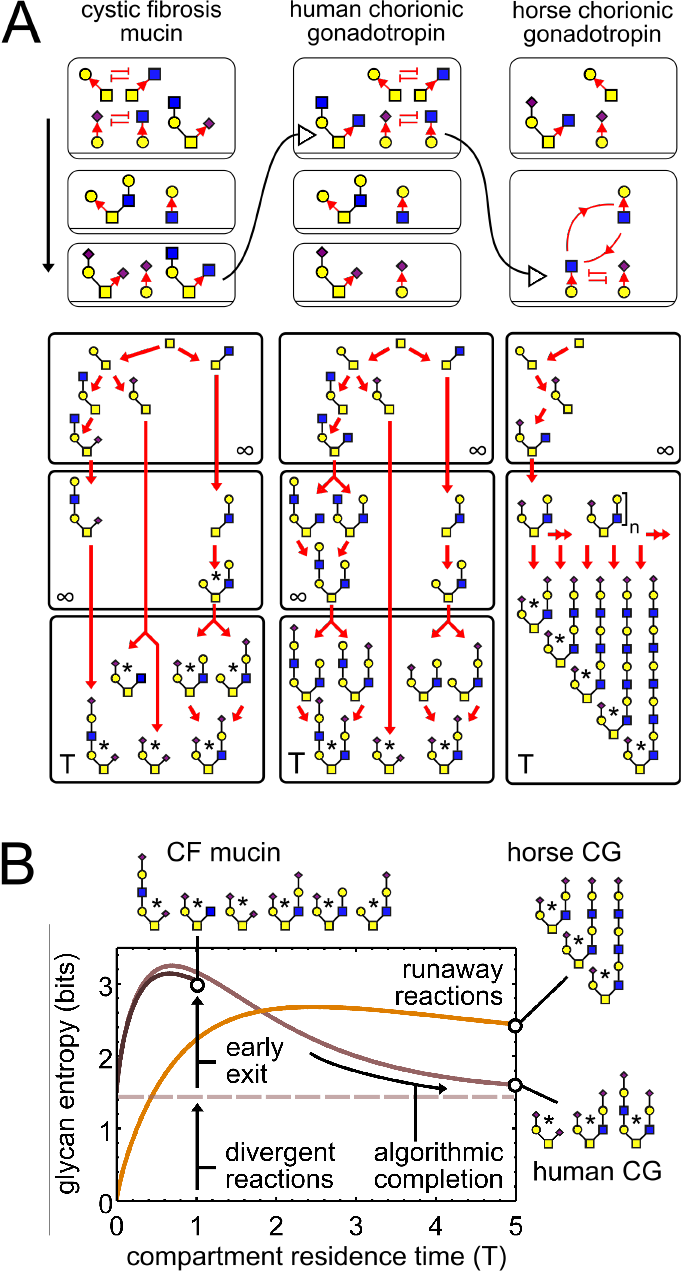
Microheterogeneity in real glycan datasets. Respiratory mucins of a cystic fibrosis patient [19], human chorionic gonadotropin from a cancer cell line [9], and horse chorionic gonadotropin [10] have distinct glycan profiles (starred oligomers; datasets from UniCarbKB [6]; we show only non-fucosylated oligomers). ***(A)*** Top: A particular localization of GTase enzymes in a multi-compartment series ordered from top to bottom, consistent with each glycan dataset. All enzymes are branch-sensitive. Blunt red arrows between two enzymes represent acceptor conflicts; cyclic arrows highlight enzymes sets that cause linkage loops. Curved black arrows highlight that the three compartmental organizations are related by a few enzyme removals or swaps. For horse chorionic gonadotropin the last two compartments are combined; this creates a runaway reaction. Bottom: Reaction networks within each compartment, starting from the same root monomer. The residence time of the last compartment in each series is *T*, so it could produce a combination of incomplete and terminal oligomers at outputs. The residence time of all other compartments is ∞, so they produce only terminal oligomers as outputs. Red arrows show single-monomer-addition reactions. Oligomers observed in each glycan dataset are starred. ***(B)*** Effect of compartment residence time on entropy of glycan profiles. We model the reaction networks in Fig. 5A as Markov processes with constant transition probabilities per unit time (PROOFS: Remark 1), and calculate the probability distribution over each possible fate of the final output oligomer. The Shannon entropy of this output distribution in bits captures the diversity of the glycan profile; approximately, it is the log-base-two of the number of distinct high-abundance oligomers. The entropy of the output distribution depends on that of the input distribution and on the residence time *T* of the final compartment. For horse CG the input entropy of the final compartment is zero, since there is a unique input oligomer; for human CG and CF mucins the input entropy is 1.5, since the input oligomers are in a 1 : 1 : 2 ratio due to divergent reactions in earlier compartments. At short residence times the entropy initially rises due to early exit of incomplete oligomers. This corresponds to the CF mucin profile. At longer residence times multiple incomplete oligomers converge to a few terminal oligomers, so the entropy decreases. This algorithmic completion corresponds to the human CG profile. The entropy continues to rise for the horse CG profile due to the synthesis of tandem repeats via runaway reactions.

The algorithmic framework also has explanatory power, providing a parsimonious account of patterns observed in existing glycan datasets. Prokaryotic glycans often contain tandem repeats [3, Chapter 21,22]. Contrast this with eukaryotic glycans: identical unbranched monomer types almost never occur in tandem (Fig. 4B); and longer tandem repeats are rare (Fig. 5). This suggests that eukaryotic cells disfavor runaway glycosylation reactions. Unlike prokaryotes, eukaryotes are able to distribute GTase enzymes across multiple Golgi compartments, thereby eliminating linkage loops. This necessarily imposes a partial order on monomer types added from root to tip on an oligomer chain (Fig. 2C). Such a partial order is evident in the characteristic layered pattern seen in many glycan oligomers, appearing as if distinct monomer types were added in pulses over time, across parallel branches. In our framework there is no need to invoke such a pulsed addition mechanism, the layering follows directly from the elimination of runaway reactions.

As anyone who has written a computer program knows, small changes to the code can result in large changes to the output. The same is true for the glycosylation algorithm: small changes in enzyme localization can result in fundamentally different glycan profiles, potentially even changing an algorithmic reaction to a runaway reaction. Real glycan datasets suggest that such changes actually do occur (Fig. 5). This has clear implications for understanding speciation, cellular identity and disease. Cells can generate distinct glycan profiles by simply re-distributing the same GTase enzymes among Golgi compartments, rather than by expressing distinct context-specific GTase enzymes in different cell types [13]. This kind of reprogramming provides a flexible way to encode cellular identity. By the same token, errors in GTase enzyme localization could be detrimental [21]. Aberrant enzyme localization [35]–[37] does cause changes in glycan profiles. The precise mechanisms by which such changes actually lead to disorders [17]–[21] are poorly understood, largely because the nature of the information stored in glycan profiles is unknown. Using the algorithmic lens to study how glycan profiles are encoded, generated and changed is the first step to understanding how cells actually read and use glycans [16].

## Methods: Definitions

### Glycan oligomer

A set of monomers linked to form a finite tree. Oligomers are grown one monomer at a time, so every oligomer or sub-oligomer (any subtree of the oligomer) has a well-defined root monomer and a well-defined direction of growth. A chain is a root-to-tip path in an oligomer. (Fig. 1A)

### Acceptor monomer

A monomer in an oligomer that can be linked to to a new donor monomer at some specific carbon, through the action of some GTase enzyme. (Fig. 1A)

### Empty acceptor monomer

An acceptor monomer with nothing linked to any carbon, except the carbon by which the monomer is linked to the oligomer. A donor monomer becomes an empty acceptor monomer once it is linked to the oligomer. (Fig. 1A)

### Branch

The entire sub-oligomer grown on a given carbon of a given monomer. If nothing is linked to a given carbon, we say the corresponding branch is empty. New branches are initiated when empty carbons are linked to donor monomers. (Fig. 1A)

### Tandem repeat

A chain in an oligomer that contains repeated instances of the same monomer type. (Fig. 1C)

### Acceptor-substrate

For branch-sensitive GTase enzymes the acceptor-substrate is a specific acceptor monomer type with an empty branch at the carbon to be linked, and specific branches or empty branches at all other carbons. For branch-insensitive GTase enzymes the acceptor-substrate is a specific acceptor monomer type with an empty branch at the carbon to be linked, regardless of other branches. (Fig. 1A,B)

### Type I acceptor conflict

One enzyme can act directly on the acceptor-substrate of another enzyme, such that the action of the first blocks the action of the second. This implies either that both enzymes compete to act on the same carbon, or that the second enzyme is branch-sensitive and acts only on the original unmodified acceptor-substrate. (Fig. 2B)

### Type II acceptor conflict

One enzyme can act on a branch of the acceptor-substrate of another enzyme, such that the action of the first blocks the action of the second. This implies that the second enzyme is branch-sensitive and acts only the original unmodified acceptor-substrate. (Fig. 2B)

### Compartment

A reaction compartment containing a set of specified GTase enzymes, and characterized by an average oligomer residence time (Fig. 1B).

### Input oligomer

An oligomer or sub-oligomer that is provided as an input to a compartment. (Fig. 1B)

### Output oligomer

An oligomer that exits the compartment after some residence time, at some stage of growth. (Fig. 1B)

### Series of compartments

An ordered set of compartments in which every output of each compartment is passed as an input to the next compartment. An initial input is provided to the first compartment, and the last compartment produces the final outputs. (Fig. 4A)

### Growth order

The order in which an oligomer is grown one monomer at a time, starting from a specific initial oligomer and leading to a specific final oligomer, through the action of successive enzymes in one or more compartments. (Fig. 2C)

### Depth-first growth order

A growth order in which, as soon as a new donor monomer is linked to an empty carbon of an acceptor monomer, the pre-existing branches of that acceptor monomer no longer grow. (Fig. 2C)

### Uniform depth-first growth

A depth-first growth order in which identical acceptor-substrates arising at any stage of growth are always linked to the same donor monomer type at the same empty carbon. This imposes a partial order on monomer types along chains, and a strict order on branch additions for each acceptor-substrate. (Fig. 2C)

### Reaction network

The nodes of a reaction network represent distinct oligomers, and its directed edges represent single-monomer-addition reactions. A reaction network shows all possible growth orders in a given compartment starting from a given input oligomer. This corresponds to all possible orders in which the growing oligomer can encounter and be acted upon by the available GTase enzymes. (Fig. 1B)

### Terminal oligomer

An oligomer with no outgoing edges in the given reaction network (labeled ^). (Fig. 1B)

### Incomplete oligomer

An oligomer with at least one outgoing edge in the given reaction network. (Fig. 1C)

### Trigger

An acceptor-substrate that cannot be fully synthesized within a compartment starting from an empty acceptor monomer input. A trigger is effectively a novel monomer type such that other donor monomers can be added to it, but it cannot be added to other acceptor monomers. (Fig. 1B)

### Linkage network

The nodes of a linkage network represent distinct monomer types or triggers; its directed edges represent acceptor-to-donor linkages at specific carbons. To construct a compartment’s linkage network we add an arrow from one monomer type to another if the corresponding acceptor-to-donor linkage occurs on any oligomer that can be synthesized starting from any empty acceptor monomer input. We need consider only oligomers whose height is less than or equal to that of the tallest acceptor-substrate of any GTase enzyme in the compartment. An enzyme whose acceptor-substrate is a trigger will not be represented among the arrows we have added so far. We must explicitly list each trigger and draw an arrow from its acceptor monomer to the relevant donor monomer type. This gives the full linkage network of the compartment. (Fig. 1B, Fig. 2A,B,D)

### Runaway reaction

A reaction network, starting from a given input oligomer, that has at least one infinite path. (Fig. 1C)

### Divergent reaction

A reaction network, starting from a given input oligomer, that has at least one fork beyond which reaction paths never reconverge. (Fig. 1C)

### Algorithmic reaction

A reaction network, starting from a given input oligomer, that has no runaway or divergent reactions and therefore converges to a unique terminal oligomer. (Fig. 1C)

### Algorithmic compartment

A compartment that has a well-defined input-to-output map: it converts each possible input to a corresponding unique output. (Fig. 1C)

### Algorithmically achievable

An input/target oligomer pair is said to be algorithmically achievable if there is a series of one or more compartments that converts the input to the target as the unique final output. (Fig. 4C)

## METHODS: PROOFS

### Remark 1.

Conditions for an algorithmic compartment.

Given a reaction network starting from some input oligomer, define a truncated network by cutting off every reaction path at an arbitrary oligomer height much larger than the input oligomer height plus the number of monomer types. The terminal oligomers of the truncated network are then either terminal oligomers of the original network, or oligomers containing arbitrary numbers of added tandem repeats. We model the truncated network as a Markov process with constant transition probabilities per unit time. The growing oligomer exits the compartment after some residence time *T*. For any finite *T* the exit probability for any oligomer in the truncated network is non-zero. As *T* → ∞ the exit probability for any incomplete oligomer tends to zero, while the exit probability for any terminal oligomer or arbitrary tandem repeat oligomer is non-zero. The compartment will be algorithmic if and only if: (a) it has no runaway reactions so there are no arbitrary tandem repeats; (b) it has no divergent reactions so every input leads to a unique terminal oligomer; and (c) it has an infinite residence time so only this unique terminal oligomer exits as an output. (Fig. 1C)

### Lemma 1.

Runaway reactions ⇔ linkage loop.

*Proof:* Consider a reaction network starting from some input oligomer. Keep all enzymes involved in this reaction network, remove other enzymes from the compartment. Suppose the reaction network has an infinite runaway path. Each reaction corresponds to the addition of one monomer to an oligomer. Therefore the reaction network contains at least one oligomer with an arbitrarily long root-to-tip chain. Since the number of monomer types is finite, the chain must include at least two instances of the same monomer type added within the compartment. Therefore the compartment’s linkage network contains a loop. Conversely suppose a compartment’s linkage network contains a loop. Then there is at least one monomer type, added at some step of the reaction network, on which a branch can be grown that includes another instance of the same monomer type. This process can be iterated ad infinitum to produce arbitrary tandem repeats. Therefore the reaction network contains an infinite runaway path. (Fig. 2A)

### Lemma 2.

Divergent reactions ⇒ acceptor conflict.

*Proof:* Consider a reaction network starting from some input oligomer. Keep all enzymes involved in this reaction network, remove other enzymes from the compartment. Suppose the reaction network has a divergent reaction. A fork in a reaction network occurs when two enzymes can act on the same oligomer. If the enzymes could act in either order the fork could immediately reconverge. Therefore there is at least one pair of enzymes such that the action of the first enzyme blocks the action of the second. There are only two ways this can occur: one enzyme acts directly on the acceptor-substrate of another (Type I acceptor conflict) or one enzyme acts on a branch of the acceptor-substrate of another (Type II acceptor conflict) (Fig. 2A,B). The converse is not true: acceptor conflicts do not imply divergent reactions, since reaction paths might later reconverge due to the action of some subsequent enzyme. (Fig. 2B)

### Lemma 3.

Every finite reaction network contains at least one growth order from the input oligomer to one of its terminal oligomers that is (a) uniform depth-first and (b) achievable in a series of single-enzyme infinite-residence-time compartments.

*Proof:* Consider a finite reaction network starting from some input oligomer. Keep all enzymes involved in this reaction network, remove other enzymes from the compartment. Since there are no infinite runaway reactions, there are no linkage loops. Every possible growth order starting from the input oligomer leads to some terminal oligomer. Any time a branch-insensitive enzyme acts at any step of any growth order, add a new branch-sensitive enzyme with the corresponding acceptor-substrate; then remove all branch-insensitive enzymes. This leaves the reaction network unchanged. Now there is at least one growth order in which a given branch-sensitive enzyme with a Type I or Type II conflict is completely blocked from acting because its acceptor-substrate (Type I) or a branch of its acceptor-substrate (Type II) is modified by some other enzyme. This growth order is retained when we remove the blocked enzyme. By iterating this procedure we eliminate every Type I and Type II conflict. The resulting compartment is algorithmic, so every growth order leads to just one of the original terminal oligomers. Every such growth order must be depth-first (there are no Type II conflicts, so every enzyme only acts on acceptor-substrates with fully extended branches) and uniform (there are no Type I conflicts and the final oligomer is terminal, so identical acceptor-substrates are identically extended). Among these growth orders there is at least one in which each given enzyme acts successively on every available instance of its acceptor-substrate on the oligomer, before the next enzyme acts. This growth order is retained even once we replace the compartment by a series of single-enzyme infinite-residence-time compartments.

### Theorem 1.

An input/target oligomer pair is algorithmically achievable in a single compartment if and only if there is a uniform depth-first growth order from the input to the target.

*Proof:* Suppose an input/target oligomer pair is algorithmically achievable in a single compartment. The reaction network starting from the given input oligomer must be algorithmic, with the target as its unique terminal oligomer. Therefore there is a uniform depth-first growth order from the input to the target (by Lemma 3). Conversely, suppose there is a uniform depth-first growth order from the input to the target. Each step of the growth order corresponds to the action of some branch-sensitive GTase enzyme. We now check what happens when just these enzymes are simultaneously present in a single compartment. Since growth is depth-first, there are no Type II conflicts. Since identical acceptor-substrates are identically extended, there are no linkage loops or Type I acceptor conflicts. Therefore the compartment contains no runaway or divergent reactions, and the final oligomer in the growth order is terminal. In particular the reaction network starting from the given input oligomer is algorithmic, with the target as its unique terminal oligomer. At infinite residence times the compartment will produce the target as the unique output oligomer starting from the given input, so the input/target oligomer pair is algorithmically achievable. (Fig. 3)

### Theorem 2.

An input/target oligomer pair is algorithmically achievable in a series of N compartments if and only if there is a growth order from the input to the target that can be fully decomposed into N uniform depth-first stretches.

*Proof:* Suppose the input/target oligomer pair is algorithmically achievable in a series of *N* compartments. Every possible growth order starting from the initial input oligomer leads to the target as the unique final output oligomer. The outputs of each compartment are passed as inputs to the next compartment. Each reaction network starting from each input of each compartment is finite (since there are no arbitrary tandem repeats) and therefore has at least one uniform depth-first growth order from its input to one of its terminal oligomers (by Lemma 3). Each compartment will produce all its terminal oligomers as outputs (potentially along with incomplete oligomers at finite residence times).

Therefore there is a growth order starting from the initial oligomer, passing via uniform depth-first stretches through just one terminal oligomer of each successive compartment, and producing the terminal oligomer of the last compartment as the final output. It also follows (by Lemma 3) that there is a series of single-enzyme infinite-residence-time compartments that achieves each depth-first stretch. Conversely, suppose there is a growth order from the input to the target that can be fully decomposed into *N* uniform depth-first stretches. Then each stretch can be achieved within a single compartment (by Theorem 1) so the input/target oligomer pair is algorithmically achievable in a series of *N* compartments. (Fig. 4)

### Corollary 2.1.

An input/target oligomer pair is algorithmically achievable if and only if there is a series of single-enzyme infinite-residence-time compartments that converts the input to the target as the unique final output.

## ACKNOWLEDGEMENTS

We thank Ajit Varki for introducing us to glycans, and Arnab Bhattacharyya for helping us view glycosylation through the algorithmic lens. We thank Ramya Purkanti, Somya Mani, Mugdha Sathe, Kabir Husain and Amit Singh for useful discussions.

## AUTHOR CONTRIBUTIONS

MT conceived the project. AJ and MT designed and carried out the analysis, proved the theorems, and wrote the paper.

